# Eco-evolutionary dynamics further weakens mutualistic interaction and coexistence under population decline

**DOI:** 10.1101/570580

**Authors:** Avril Weinbach, Nicolas Loeuille, Rudolf P. Rohr

## Abstract

With current environmental changes, evolution can rescue declining populations, but what happens to their interacting species? Mutualistic interactions can help species sustain each other when their environment worsens. However, mutualism is often costly to maintain, and evolution might counter-select it when not profitable enough. We investigate how evolution of mutualism affects the coexistence of two mutualistic species, e.g. a plant-pollinator or plant-fungi system. Specifically, using eco-evolutionary dynamics, we study the evolution of the focal species investment in the mutualistic interaction of a focal species (e.g. plant attractiveness via flower or nectar production for pollinators or carbon exudate for mycorrhizal fungi), and how it is affected by the decline of the partner population with which it is interacting. We assume an allocation trade-off so that investment in the mutualistic interaction reduces the species intrinsic growth rate. First, we investigate how evolution changes species persistence, biomass production, and the intensity of the mutualistic interaction. We show that concave trade-offs allow evolutionary convergence to stable coexistence. We next assume an external disturbance that decreases the partner population by lowering its intrinsic growth rate. Such declines result in the evolution of lower investment of the focal species in the mutualistic interaction, which eventually leads to the extinction of the partner species. With asymmetric mutualism favouring the partner, the evolutionary disappearance of the mutualistic interaction is delayed. Our results suggest that evolution may account for the current collapse of some mutualistic system like plant-pollinator ones, and that restoration attempts should be enforced early enough to prevent potential negative effects driven by evolution.

## Introduction

Facing current global change, evolutionary mechanisms can help maintain biodiversity. Evolutionary rescue [1,2] corresponds to the selection of new traits in population collapsing with environmental changes, that allow for a demographic bounce. The signature for evolutionary rescue is the increase in frequency of the allele and corresponding phenotype robust to the new environment, correlatively to the population bounce. While this can be easily highlighted in lab experiments, it has so far been seldom observed in nature. In their review Carlson and collaborators [2] cite, for example, a previous study showing the adaptation of some *Chlorella* species, but not all, after strong acidification of many Canadian lakes with industrial pollution.

However, species are not isolated from one another and interaction might interfere with this evolutionary process. Mutualism is an interaction that has already been intensively studied, and proven to be fragile to Global Changes [3]. Echoing to that loss of interactions is that of biodiversity and system services such as pollination [4] and seed dispersal [3] and effective carbon and nutrient cycles [6]. While several reviews like that of Potts and collaborators [7] point out the critical ecological crises we are undergoing, Toby Kiers and collaborators [3] add that mutualism, by binding species to a common fate, could create an evolutionary breakdown. With environmental changes, mutualism can become costly to maintain. Aside from coextinction of the two interacting species, evolution can lead to mutualism loss, partner switch, or even a shift to antagonism. They thoroughly present how the different types of mutualisms are specifically sensitive or resistant to breakdown depending on the global change drivers. For example, plant-pollinator mutualism could be strongly affected by climate change and habitat fragmentation, while plant-rhizosphere mutualisms will be more affected by nutrient enrichment and the introduction of exotic species.

Mutualistic systems, like all other systems of interacting species, will respond differently to global change [8]. One species can be severely impacted by environmental disturbances, showing a strong decrease in its density, while its mutualistic partner species might have higher standing genetic variabilities or larger population sizes, so that it could be presumed to adapt. However, because the fitness of the two partners are positively linked (mutualistic interaction), the fitness decrease of the first species may eventually harm the second, reducing its potential for evolutionary rescue and slowing its evolution (evolutionary inertia, [9]). For mutualism, and especially obligatory ones, this might lead to species extinction, driven by the evolutionary disinvestment of its interactor. This effect is called an evolutionary murder (name suggested by [10]).

For instance, plants have been shown to evolve rapidly to changing pollinator populations [11–14]. A recent study from Gervasi and Schiestl [15] experimentally showed that changes in pollinator communities affect plant trait evolution after only eleven generations. Exposed to bumblebees, which are very efficient pollinators of *Brassica rapa*, the plants evolved toward more attractive traits to those pollinators (e.g. traits attracting pollinators such as volatile organic compounds, flower size, or plant height). Moreover, hoverflies, which are less efficient pollinators of *B. rapa*, caused a 15-fold increase in self-reproduction and a reduction in plant attractiveness. Given these experimental results, the current change and reshaping of pollinator communities may affect the evolution of plant species, which in turn could influence coexistence with their interacting pollinators, i.e., an eco-evolutionary feedback loop.

Plant-mycorrhizal fungi interaction is another type of mutualism affected by global changes. The mycorrhizal fungi can for example fix inorganic nitrogen and provides this essential nutrient to the plant, who, in exchange, transfers via its roots carbon products to the fungi. Several experiments already showed that enriching the soil in nitrogen (often from anthropogenic sources in natural environments) disturbs this mutualistic exchange, by inducing a shift in the allocation to the mycorrhizal structures [16] and the composition of the mycorrhizal community [17]. This can in turn affects the plant community, and can even facilitate the invasion of alien plant species [18].

Theoretical studies have investigated the ecological [19–22] and evolutionary dynamics [3,23–26] of mutualistic communities such as plant-pollinators or plant-fungi. In particular, the evolution of plant selfing with changing pollinator communities has been studied in several papers [27–29]. Thomann and collaborators [30] even suggested that the decrease in pollinator richness and density could intensify pollen limitation. They propose that plants could in turn adapt either by increasing autonomous selfing or reinforcing the interaction with pollinators. Here we study the consequences of a declining population (e.g. pollinator collapse) on the eco-evolutionary dynamics of a two-species mutualistic system (e.g. plantpollinator or plant-fungi). Specifically, we study the eco-evolution of the investment in mutualism of a focal species. To do so, we use the adaptive dynamics framework. This framework explicitly accounts for the eco-evolutionary feedback loop between the two species. We clarify when evolution leads to high or low investment in mutualism and determine the conditions under which evolution leads to the coexistence of the whole system. We then show that a declining partner population often results in a counter-selection of the investment in mutualism of the focal species, which eventually enhances the population declines. For simplicity in the narrative, in the following, we will use the example of a plant-pollinator system. The adaptive trait is the plant investment, and the declining population is the pollinator. However, our approach remains general and can be applied to other mutualistic systems.

## Plant-pollinator model and ecological dynamics

We consider a simple system with two interacting species; a plant with biomass density, and a pollinator with biomass density. Note that this model is formulated as a general model of mutualism rather than very specifically tied to plant-pollinator interactions so that results may also concern other mutualistic systems. The community dynamics are given by a Lotka-Volterra type model:

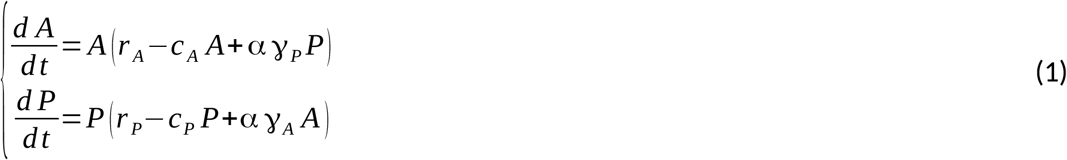

A schematic view of the system is given in figure 1. The parameters *r_A_* and *r_P_* correspond to the intrinsic growth rate of the pollinator and plant populations, respectively. We assume *r_P_* to be strictly positive because of other reproduction means, e.g. vegetative reproduction or autogamy. The intrinsic growth rate of the pollinator (*r_A_*) can be positive (e.g., interaction with other plants) or negative. Parameters *C_A_* and *C_P_* modulates intraspecific competition for the two species. Mutualistic interactions are given by *αγ_A_* and *αγ_P_*, with *γ_P_* the energetic gain provided by the plant (via nectar, pollen and/or other plant exudates) to the pollinator, and *γ_A_* the fertilisation provided by the pollinator to the plant. Because we consider mutualism as the net benefice obtained by both species, both *γ* parameter values are assumed positive in our model. We modulate the intensity of the interaction between the two species with the parameter α. While the interaction depends on biological traits from both interactors (e.g. pollinator morphology or flight capacities, plant attractiveness), we have chosen to model it as a plant-dependant trait and have therefore linked it to other plant traits via a trade-off function (figure 2). We interpret it here as the attractiveness of the plant for the pollinator, and it corresponds to the trait that is under selection in the rest of the study. This plant attractiveness includes investment in various characters such as the number of flowers, their shape, their colour, volatile organic compound (VOCs) that attract insects with their odour, plant height, flowering duration or nectar quantity and quality (see part II in Willmer (2011) [28]).

**Figure 1:**
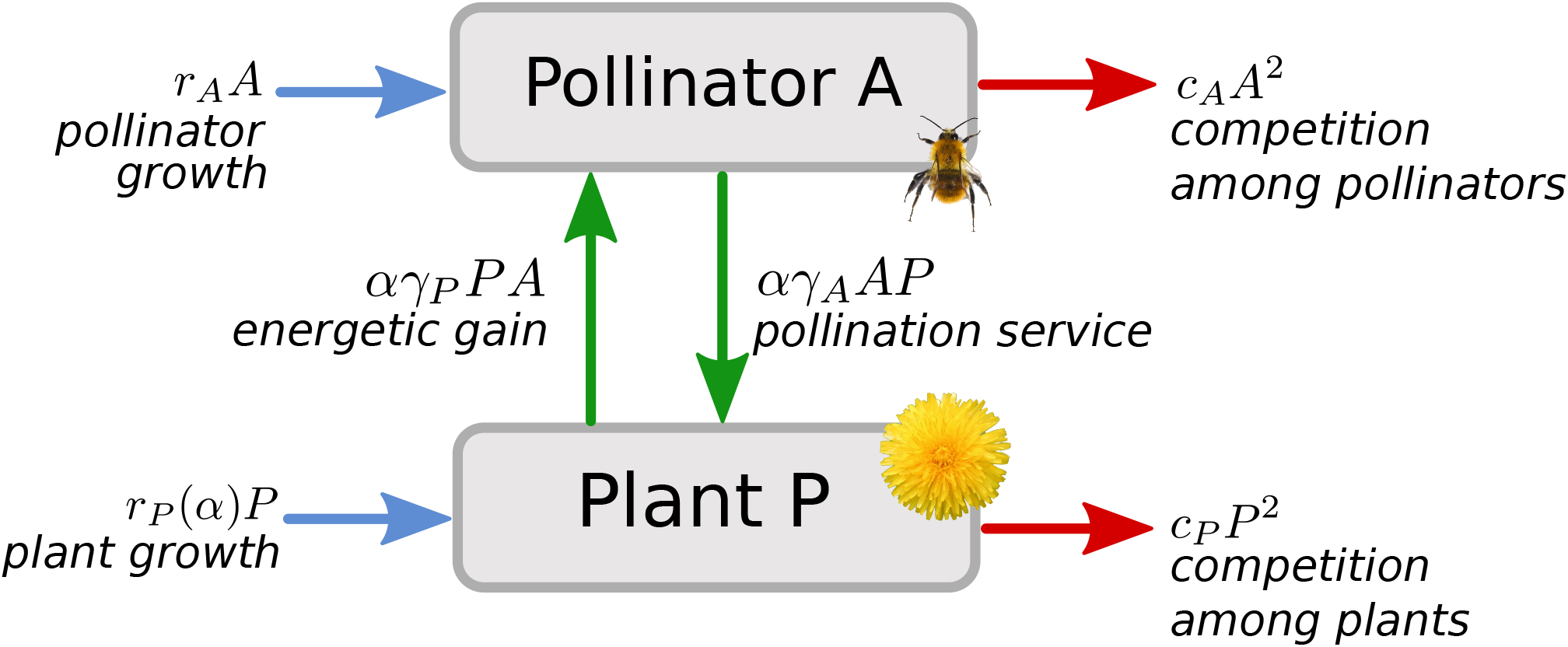
Population variation rates of plant and pollinator. Blue arrows indicate the density variations via other means than the mutualistic interaction, green arrows the effects of the mutualistic interaction, and red arrows the effects of intraspecific competition. Note that the plant intrinsic growth rate r_P_ is in trade-off with the plant attractiveness α. The parameters are described in the main text. B).

**Figure 2:**
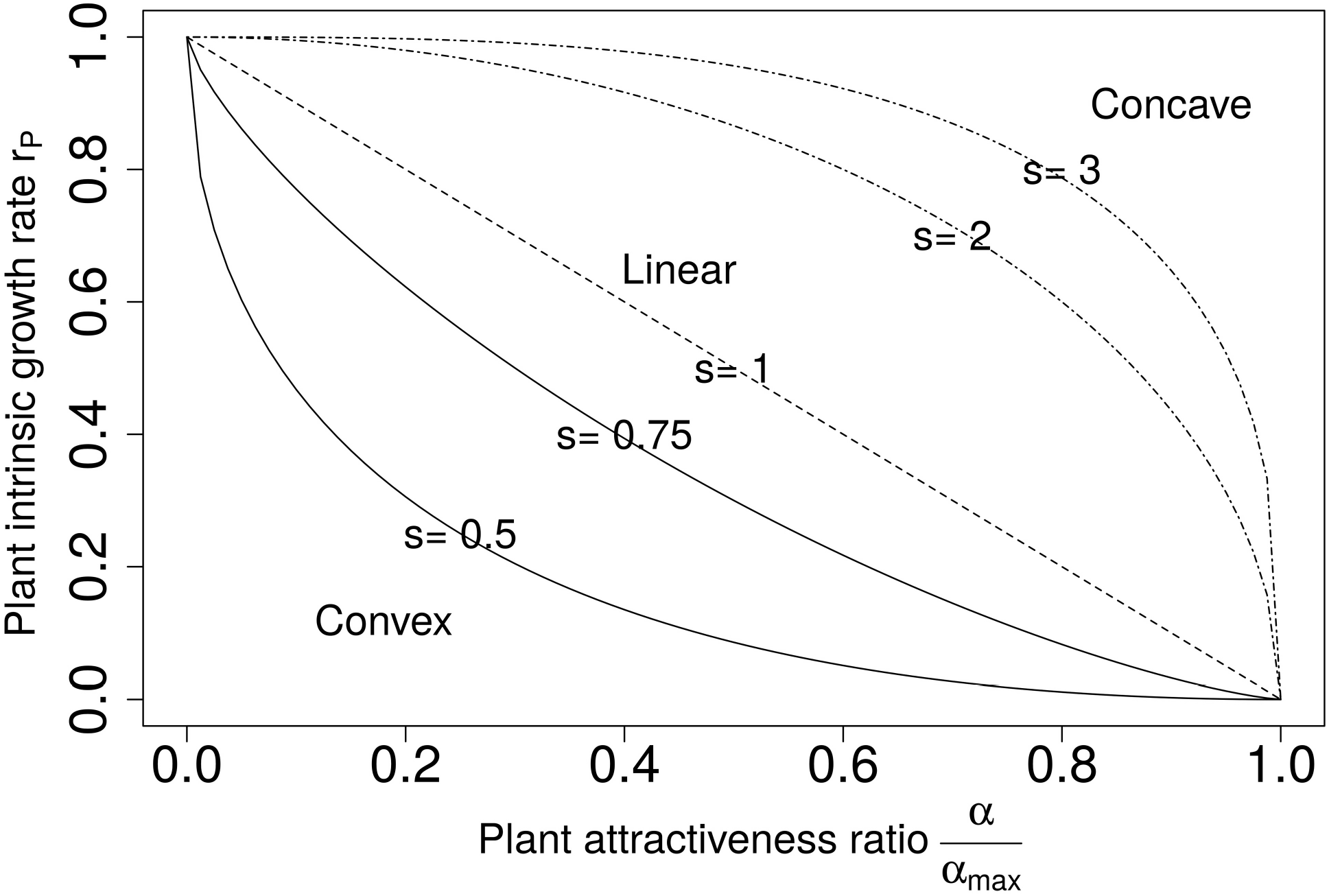
Variation of the attractiveness ratio 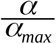 with the plant intrinsic growth rate r_P_ depending on the trade-off strength. Continuous lines show convex trade-offs, the dashed line a linear trade-off, and dashed-dotted lines concave trade-offs.

Extrapolating from previous results [22], coexistence is stable provided:

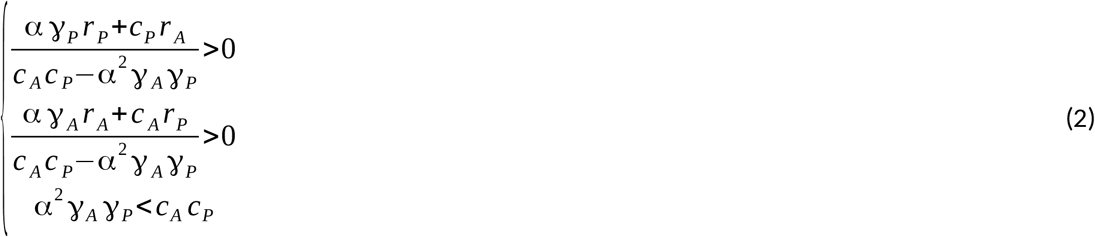

The first two inequalities give the condition for the existence of an equilibrium point allowing positive densities (i.e. feasibility conditions). The last inequality ensures the stability of the equilibrium. According to Goh [31], in the case of two interacting species, conditions for a feasible and locally stable equilibrium with intraspecific interactions regulating the population densities implies its global stability. The globally stable equilibrium is then:

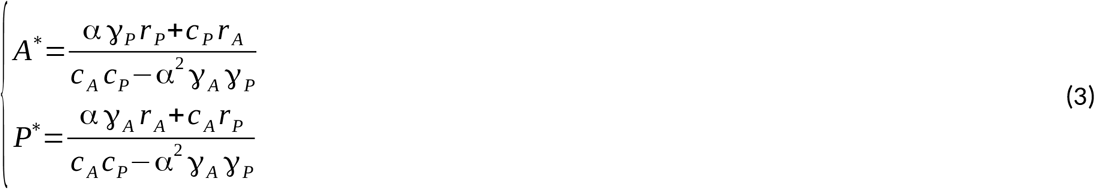

If the stability condition is not fulfilled, i.e., interspecific mutualism is stronger than intraspecific competition, the positive feedback loop resulting from interspecific mutualism may drive the system towards infinite growth. In such cases, other limiting factors (e.g. pathogen, predators, or new competitors) eventually regulate the populations. Since these factors are not taken into account in our model assumptions, we define a maximum plant attractiveness *α_cl_* below which stability is warranted:

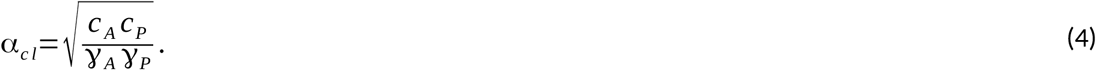

We allow the evolution of α between zero (no investment in attractiveness) and this maximal level α_*max*_ < α_*cl*_. We could also have controlled the infinite growth of our system by choosing a saturating function for the mutualistic interaction (Holling type II or other, e.g. [32]). Our choice of a linear functional response, however, allows explicit analytical computations and has the advantage to keep the model general and applicable to mutualistic interactions other than pollination, i.e. ant-plant, plant-rhizosphere or coral-zooxanthellae mutualisms [3].

## Evolution of plant attractiveness

We study the evolution of plant attractiveness (α), assuming an allocation trade-off affecting the plant intrinsic growth rate *r_P_* [33]. Its biomass can grow either via a reproduction process dependent on the interaction with its mutualist (e.g. pollination) whose intensity is controlled by its attractiveness α, or via intrinsic growth (e.g. vegetative growth) and self-reproduction. The plant has a given quantity of energy that is divided between these two growth modes [33,34], so that we assume *r_P_* to be a decreasing function of the attractiveness α :

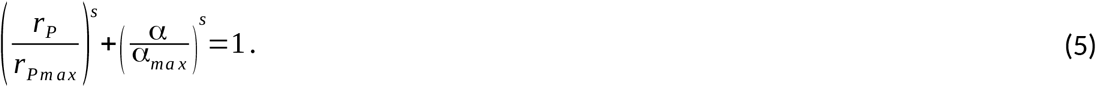

The plant maximal intrinsic growth rate *r_Pmax_* can be fixed to one without loss of generality, by rescaling time unit:

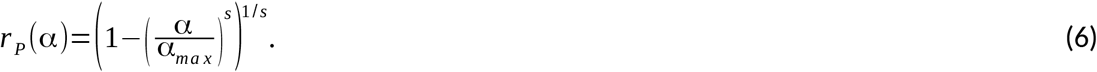

The *s* controls the trade-off shape. When *s*=1 there is a linear relationship between *r_P_* and α. When 0<s<1 the trade-off is convex. On the opposite, *s* >1 produces a concave trade-off (as shown in figure 2).

We follow the evolution of plant attractiveness using adaptive dynamics techniques [32,33]. Under adaptive dynamics hypotheses (see supplementary material section A for a full description of the method and hypotheses) we can model the evolution of plan attractiveness and its consequences on species density dynamics, and the feedback of species density on the evolutionary process [35]. Evolution then proceeds by the successive invasions and replacements of resident by mutant populations. Such dynamics are approximated, given rare and small mutations, by the canonical equation [35]:

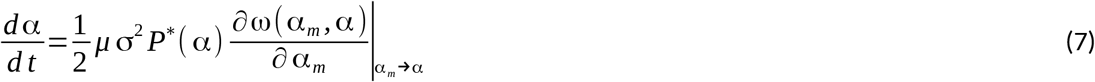

The term 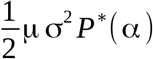 encapsulates the phenotypic variability brought by the mutation process on which selection can act. The last term, called the selective gradient, is based on the variations of the relative fitness of mutants α_*m*_ given a resident α. It gives the direction of evolution; a positive gradient selects larger attractiveness, while a negative gradient selects smaller trait values. The relative fitness of the mutant is computed as the *per capita* growth rate of a rare mutant population in a resident population at ecological equilibrium (3):

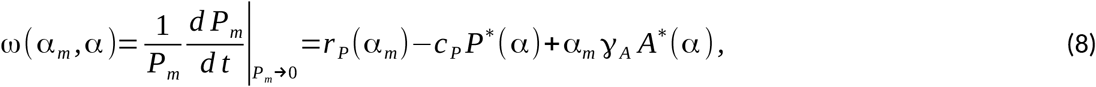

Eco-evolutionary equilibrium (called a singular strategy) occurs when the phenotypic trait stops varying, i.e. equation 7 equals 0. Since its first part is always positive, it corresponds to 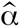 values for which the selective gradient is null:

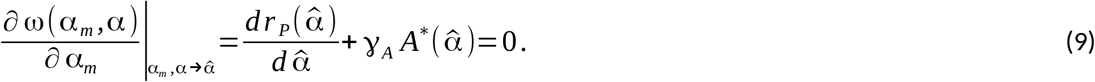

At singularities, costs in terms of energy dedicated to alternative means of reproduction 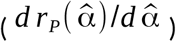 therefore match pollination benefits 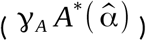. The existence of a singular strategy is not enough to guarantee that evolutionary dynamics locally lead to it (convergence condition) or that it persists (non-invasibility condition, i.e. resistance to invasion by nearby mutants). A singular strategy that is both convergent and non-invasible is called a continuously stable strategy (CSS) [37]. Evolution toward a CSS guarantees the coexistence of the two species. This and other singularity types are presented in figure 3. Calculation of the second and cross-derivative of the fitness function determines criteria for convergence and invasibility [38]. The mathematical computation for the existence of singular strategies and their convergence and invasibility properties are detailed in the supplementary material sections A and B.

**Figure 3:**
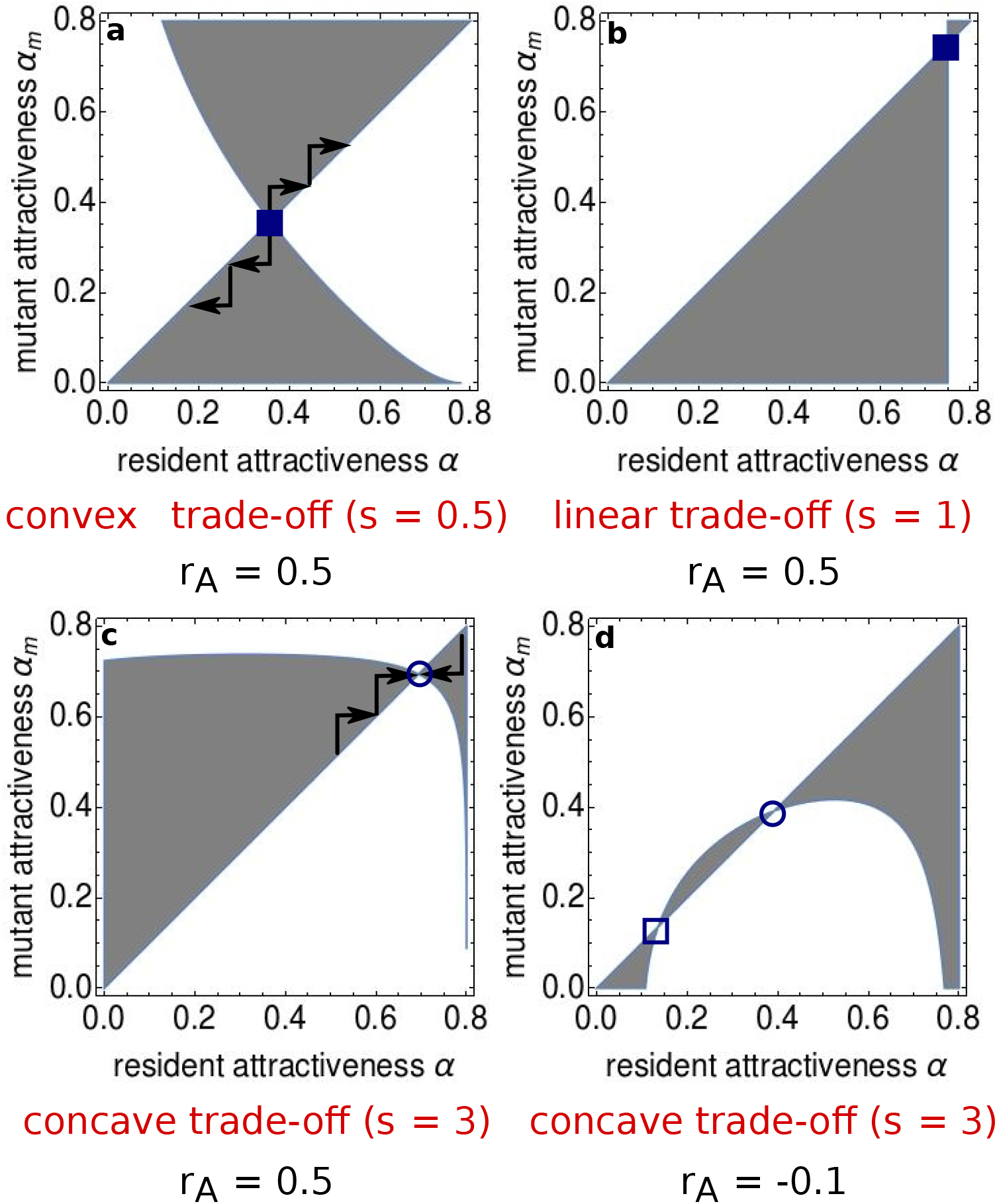
Pairwise invasibility plots (PIPs) representing the invasibility potential of a rare mutant within a resident plant population at ecological equilibrium. Grey areas indicate that the mutant relative fitness ω(α_m_, α) is positive, so that it invades and replaces the resident population. In panels a and c, arrows show the direction of evolutionary trajectories. The system exhibits several singular strategies depending on the parameter values. Circles represent convergent strategies, whereas squares are non-convergent. Filled symbols represent invasible strategy, while not filled symbols are non-invasible. In panels a and b, the singular strategy is non-convergent and invasible (repellor). In panel c, the singular strategy is convergent and non-invasible (CSS). Panel d displays two strategies, one CSS and one which is non-convergent and non-invasible (Garden of Eden). Parameter values are: c_A_=c_P_=γ_A_=γ_P_=1, and α_max_= 0.8 * α_cl_.

Equation (9) can be solved analytically for particular sets of parameters (e.g. in the linear case when *s*=1, see supplementary material section A). For other cases, we graphically determine convergence and invasibility using pairwise invasibility plots (figure 3). It is possible to show (supplementary material section B), as illustrated in figure 2, that among the particular trade-offs that we study (equation 6), only concave allocation trade-offs lead to non-invasible strategies. Therefore, CSSs, being non-invasibles, are only obtained with a concave trade-off function. Convergence depends on the pollinator’s intrinsic growth rate (figure 3c and 3d). Mathematical analyses show that linear trade-offs lead to singular strategies that are not convergent (supplementary material section B). In that specific case we can explicit the formula of the attractiveness value at eco-evolutionary equilibrium (equation (A7) in supplementary material section A). We observe that the plant investment in attractiveness increases when plant or pollinator intrinsic losses as high or pollinator intrinsic growth is low. In that case an increase in the attractiveness value at eco-evolutionary equilibrium compensate for these lower intrinsic gains. For non-linear trade-offs, convergence criteria cannot be solved and we rely on numerical investigations and PIPs.

For positive pollinator intrinsic growth rate, given concave trade-offs, we obtain only one convergent stable singular strategies (CSS) at which ecological coexistence is granted. For negative pollinator intrinsic growth rate, the system exhibits a second singular strategy that is a Garden of Eden (non-convergent and non-invasible), i.e. a stable strategy that can never be reached by nearby mutants. While the conditions of existence of multiple singularities cannot be completely mathematically derived, our results suggest that it occurs for very concave trade-offs. The case were pollinator growth rate entirely relies on the mutualistic interaction (*r_A_* = 0) can for instance be analysed mathematically and reveals that two singularities will emerge when s>=2 (supplementary material section C).

For convex trade-offs (figure 3a and 3b), we always observe repellors (non-convergent and invasible). Starting above the repellor, attractiveness increases to reach the maximum value (α = α_*max*_) and the plant growth relies only on the mutualistic interaction. In that case plant pollination can only be maintained if pollinators are present and with a positive intrinsic growth rate (obligatory mutualism on the plant side). Starting below the repellor, attractiveness evolves to zero, so that the two species no longer interact at the end of the evolutionary dynamics (e.g. complete selfing or clonal reproduction). As there is no more interaction it is trivial that pollinators are maintained only if their intrinsic growth rate is positive. In the following, because we are interested in the species long-term coexistence with intermediate investment in the mutualistic interaction (ie, CSS singularities), we only study concave trade-off functions (i.e. *s*>1).

## Consequences of pollinator population decline

Now that we have characterised the eco-evolutionary dynamics of the plant-pollinator system, we study how pollinator decline may affect its outcome. We simulate less favourable environmental conditions for pollinators (e.g. habitat fragmentation, pesticides, diseases) by decreasing their intrinsic growth rate (*r_A_*). We illustrate the effects of this disturbance through Ecology-Evolution-Environment (*E*^3^) diagrams [1,39]. These diagrams, presented in figures 4 and 5, show the outcome of eco-evolutionary dynamics as a function of the environmental parameter, here the pollinator intrinsic growth rate *r_A_*. Figure 4 exhibits two types of singular strategies depending on the pollinator intrinsic growth rate.

**Figure 4:**
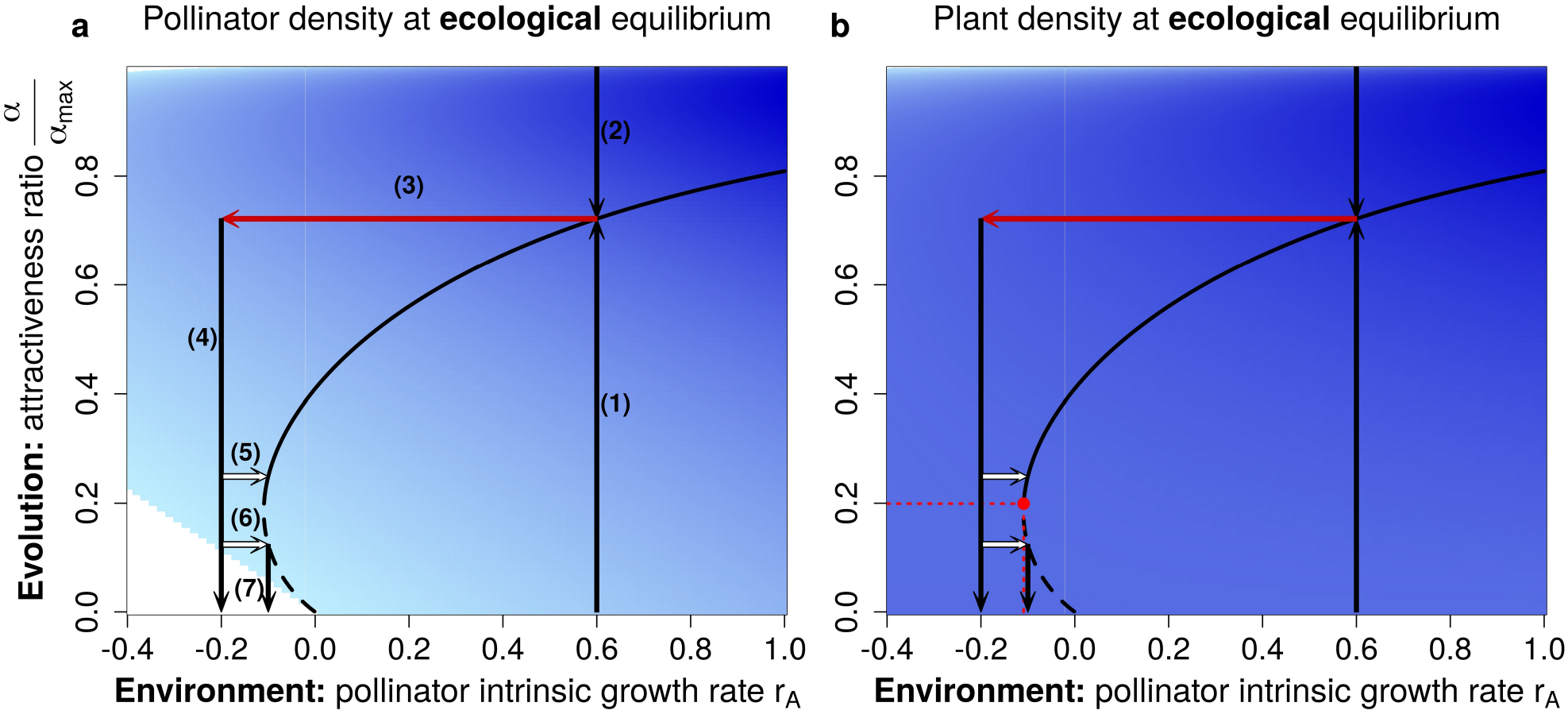
Ecology-evolution-environment (E^3^) diagram representing the impact of pollinator environmental deterioration on the evolution of plant attractiveness and on pollinator (panel a) and plant (panel b) equilibrium biomass densities. White areas show parameters for which extinction occurs for either plants or pollinators. The blue intensity correlates with population densities of pollinators (panel a) or plants (panel b). Black lines show the position of singular strategies; continuous lines show convergent and non-invasible singular strategies (CSS), and dashed lines show Garden of Edens (non-invasible, divergent). Vertical black arrows (1, 2, 4, 7) display the direction of evolution. Environmental disturbance is represented by a red arrow (3). White arrows (5, 6) represent restoration attempts at different times along the evolutionary trajectory. On panel b) the red point and dotted lines represent the lowest r_A_ and 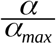 values for allowing a CSS, therefore the maintenance of the mutualistic interaction. This point is what we call an eco-evolutionary tipping point. Parameters values are s=2.5, c_A_=c_P_ = γ_P_= 1, γ=0.2, and α_max_=0.8*α_cl_. Similar E^3^ diagrams can be found in [1,39].

**Figure 5:**
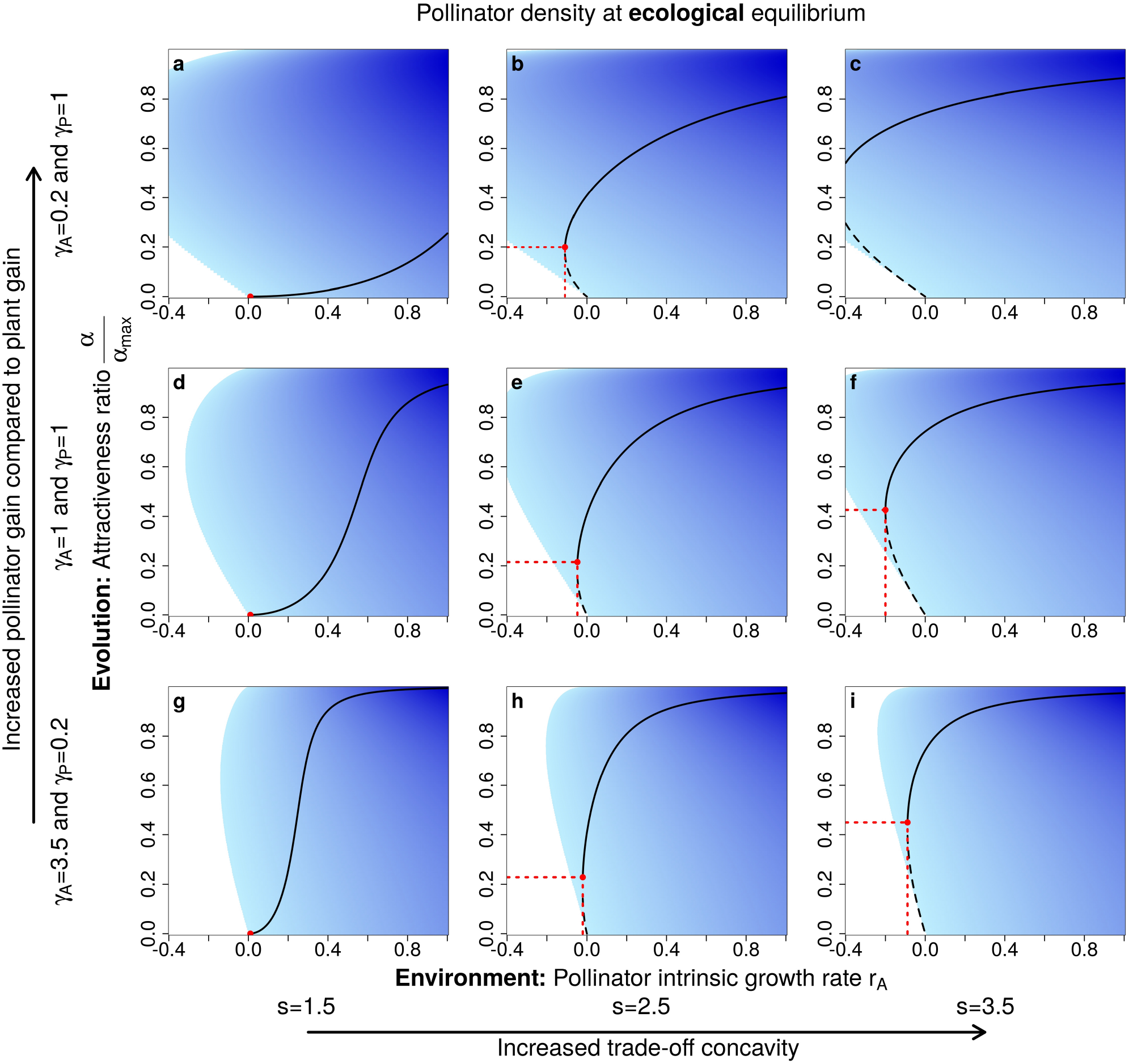
Influence of trade-off shape and mutualistic gains on diagrams. Columns differ in tradeoff concavity. Lines differ in the asymmetry of mutualistic gains: in the top line (panels a,b, and c) pollinators benefit more than plants; the middle line (panels d,e, and f) shows equal gains while in the bottom line plant gains are larger. Red point and dotted lines represent the lowest r_A_ and 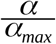 values for allowing a CSS allowing the maintenance of the mutualistic interaction. Colours and lines are the same as in figure 4. The parameter values are c_A_=c_P_=1 and α_max_=0.8*α_cl_.

For positive pollinator intrinsic growth rates (*r_A_*>0), i.e. in “good pollinator environments”, we observe a convergent and stable singular strategy (CSS, continuous line). Any ecological system with positive *r_A_* evolutionarily converges toward intermediate attractiveness α (arrows (1) and (2) in figure 4a).

For negative pollinator intrinsic growth rate *r_A_* < 0, i.e. in “bad pollinator environments”, we also find a none convergent strategy, a Garden of Eden (GOE, dashed line). In this case, the system exhibits evolutionary alternative stable states. When plant attractiveness is above the Garden of Eden value, evolution converges toward the CSS, while when below the GOE value, selection leads to ever-decreasing attractiveness that eventually leads to the disappearance of the mutualistic interaction.

If we consider environmental degradation, i.e. a strong decrease of *r_A_* (red arrow (3) in figure 4a), in the absence of evolution, both plant and pollinator populations have positive biomass densities at ecological equilibrium (blue backgrounds on figure 4a and b). However, considering evolution, plant attractiveness is counter selected as pollinators are too rare to compensate for the intrinsic costs of attractiveness. Eventually, evolution drives pollinator populations to extinction; an evolutionary murder depicted by arrow (4) in figure 4a. Faced with the crash of pollinator populations, restoration attempts may be undertaken (i.e. an increase in *r_A_* value, e.g. by suppressing pesticides or adding other plant resources for pollinators). Early intervention, depicted by arrow (5), can restore a stable mutualistic interaction. Delayed restoration attempts (white arrow (6)), do not allow such a rescue, as evolutionary trajectories will counteract their effects and lead to the extinction of the pollinator (arrow (7)). Note that here we separate timescales for simplicity, and consider that deterioration and restoration are fast compared to the evolutionary dynamics, hence horizontal arrows for these environmental changes.

Finally, we study how trade-off shapes and asymmetry of mutualistic gains affect the eco-evolutionary outcome (figure 5). With increasing concavity, the minimum value of pollinator intrinsic growth rate *r_A_* that allows for a CSS decreases (red dots on figure 5), increasing the coexistence domain (i.e. interval of *r_A_* values that allows species coexistence at an intermediate interaction level). More concave trade-offs therefore allow a larger coexistence domain up to negative values of *r_A_* (figure 5b, c, e, f, h, i.). For less concave trade-offs, s < 2, only a positive pollinator intrinsic growth rate *r_A_* allows coexistence (figure 5a, d, g). Negative pollinator intrinsic growth rates lead to small benefits for the plant, so that attractiveness is counter selected, eventually leading to the pollinator extinction. For stronger concave trade-offs, s > 2, (figure 5b, c, e, f, h, i) we observe qualitatively the same dynamics as in figure 4. For those trade-offs, asymmetric mutualistic gains favouring pollinators allow a larger range of disturbance, including negative intrinsic growth rates *r_A_*, before attractiveness is counterselected and extinction occurs. Therefore, an increased mutualistic gain of the pollinator relative to that of the plant facilitates the long-term coexistence of the plant-pollinator system. This produces a more robust system that eases a potential restoration process. Note, however, that favouring pollinators gain over plants leads to lower selected levels of attractiveness.(compare figure 5 a, b, c vs g, h, i). The same figure but with the plant density at ecological equilibrium can be found in the supplementary material section D. Apart from very high values of investment in attractiveness with strongly negative pollinator growth rates, plant density is always positive.

## Discussion

While from a one species perspective, evolution can help to avoid extinction by fostering adaptation and restoring positive growth rates (evolutionary rescue), we here show an example in which accounting for mutualistic interactions largely modifies this optimistic view. Here, the evolution of one species in response to disturbances acting on its mutualistic interactor selects a further decrease in the interactor population, eventually leading to the demise of the mutualistic interaction. This shows that evolution within mutualistic systems can actually be detrimental to the system’s persistence and could undermine restoration attempts. Because we have used a general model of mutualism, this mechanism may concern various systems. This clearly suggests that when investigating the impact of global changes, we need to account for eco-evolutionary dynamics of the species and their interactors.

The model we use is voluntarily simple to allow a more complete mathematical study of eco-evolutionary dynamics and to highlight the role of key *parameters* (e.g. trade-off shapes or mutualistic gains). However, it may be linked to other models that study various types of mutualism. For instance considering pollination systems and plant reproduction, in line with the presentation of the results, our model recalls previous theoretical works on plant evolution, that detail furthers the reproductive implications (e.g. [27–29]). For instance, Lepers *et al*. [28] explicitly modelled the evolution of a plant reproduction system by taking prior selfing and inbreeding depression into account. In particular, they showed that evolution toward high prior selfing (for us of lesser attractiveness) leads first to pollinator extinction (our evolutionary murder). Because they also model the cost resulting from the inbreeding depression, they show that this evolutionary murder may further lead to the extinction of the plant population. However, the model we propose may also be adapted to consider other mutualistic systems. For instance, in plant-mycorrhizae interactions, a resource exchange takes place, where plants provide carbon-based resources (e.g., sugars) to mycorrhizae while they get nutrients from the interaction. Such a situation does fit our hypotheses. The trait *α* would then be the quantity of resource provided by the plant (ie, its investment in the mutualistic interaction), and this production diverts resources from growth and reproduction, therefore fitting our trade-off hypotheses. As such, our model recalls the results of a study by de Mazancourt & Schwartz [24]. They show that mutualism can arise and be evolutionarily selected from a two-species competing model by including trading. Each species can trade the resource it extracts in excess with the other. In our system, this trading would correspond to the benefit *αγ* provided by the mutualistic interaction. Depending on the resource availability (the intrinsic growths in our model) the plant can either perform better on its own (possibly at the detriment of the fungi, as in our model) or can benefit from the mutualistic association with a mycorrhizal fungus. The mutualistic trading interaction can extend the coexistence boundaries, i.e. the resource space the two species can live in.

We are aware that the linear Lotka-Volterra structure of our model and the adaptive dynamics methods impose specific working hypotheses that can constrain the applicability of our model (e.g. ecological equilibrium between small mutation steps, asexual reproduction, panmixia). Our model is a better fit for specific reproduction types like that of geitonogamous species. Models that put into equations specific types of mutualistic interactions can better explicit the different biological processes at stake and have a more realistic view of the mechanisms, e.g. Fishman and Hadany pollination model [40]. They show that a complex and biologically detailed resource trade mutualism can be approximated by a Beddington-DeAngelis formula for trophic interactions. However, the previously cited more complex models [24,27–29] find a disinvestments in mutualism with declining efficiency similar to the one observed in our simple and more general model, in coherence with the results from the experimental evolution studies [14,15]. They can better detail the potential consequences of this disinvestment on the interactors for a specific mutualistic type.

Our results also highlight that mutualistic interactions could be more or less vulnerable to environmental changes and population declines. For instance, here, only concave allocation trade-offs between plant intrinsic growth rate and investment in mutualism lead to the maintenance of the mutualistic interaction. These trade-offs favour intermediate investments in the mutualistic interaction, while in the case of convex trade-offs, either complete investment or no investment is eventually selected, depending on initial conditions. We kept our study general because trade-off shapes are extremely difficult to measure *in vivo*, and can vary deeply depending on the environment or the species types [41].

Bistability and critical transitions have been highlighted in a variety of ecological situations (e.g. [42,43] in mutualistic system), and result from a strong positive feedback loop. Here we have a similar phenomenon but on an eco-evolutionary scale. If the evolved investment in mutualism before environment deterioration is above a certain threshold, evolution reinforces the interaction, by increasing the attractiveness values, eventually leading to a stable, coexisting system. On an ecological scale, this interaction reinforcement increases the abundance of both species, which in turn favours the evolution of the focal species investment toward higher value. Below a critical level of evolved investment, the population of the mutualistic partner species is low. Evolution then further decreases investment in mutualism, eventually leading to complete disinvestment in the mutualistic interaction. This runaway selection for decreased investment leads in our case to the evolutionary murder of the partner population by the evolving species [44]. Note that the tradeoff shape modulates the strength of the positive feedback loop. More concave trade-offs decrease the threshold value above which interaction is maintained, thereby facilitating the persistence of the system. Such dynamics have important implications. For instance, consider pollination as the mutualistic interaction. Current data suggest large decreases in pollinator abundances [45]. Such pollinator declines are often considered to be directly linked to environmental changes (e.g. habitat change, pesticides [7]). However, our results suggest that evolutionary components may also be present. If these declines favour plant strategies that offer less resource, plant evolution may enhance the observed declines. In line with these predictions, empirical observations suggest a decline of flower resources parallel to the pollinator decline [45].

On a management side, alternative stable states and critical transitions have large implications, as systems may then shift abruptly, and large restorations are needed to recover previous states [43]. The eco-evolutionary alternative stable states we describe here have similar implications. Restoration can either be a reduction of the mortality causes of the declining species (banning pesticide, ploughing controlling pests and predators) or the increase in their alternative resource source (plant sowing or nutrient addition). Here we consider that restorations are faster than evolutionary timescales. Evolution can however act fast [15], while restoration timescales largely vary from a few months (e.g., sowing high reward plants) to much longer timescales. Changes in pesticide regulations and applying these regulations may require national or international consensus. Similarly, while a change in the agricultural mode does occur (e.g. from intensive to agroecology), its dynamics happen over decades, while the evolution of plant reward may happen in just a few generations [15]. Note that, were we to consider longer restoration attempts we would still observe eco-evolutionary tipping points in our system. Such tipping points also make restoration attempts more difficult from two different points of view. First, the timing of the attempt becomes important. Restoration is only successful when achieved before the threshold attractiveness is evolved. Second, if the system becomes degraded, a small restoration attempt may not be sufficient to recover large populations, but large efforts will have to be undertaken.

While in the face of current changes in the environment, evolution can play a key role in restoring populations and maintaining diversity, our results suggest that in the case of mutualistic interactions, evolution may also favour strategies that eventually further threatens species coexistence. As such, our model echoes recent analyses that highlight the evolutionary fragility of mutualisms, given current changes [3,9]. Because our model is voluntarily simple, restricted in its number of interaction types and species, we expect evolutionary effects in complex ecological networks to be more complex and context-dependent. However, we expect that accounting for these covariations of evolutionary dynamics and changes in ecological interactions will be important, and that the effects of evolution will then not systematically be positive.

## Data, scripts and code accessibility

Scripts and codes used to produce the data and figure 2, 4, 5, and supplementary figure C1 and D1 are available online: https://doi.org/10.5281/zenodo.5552426.

## Acknowledgements

NL and AW were supported by the French National Research Agency (ANR) through project ARSENIC (grant no. 14-CE02-0012). AW was additionally supported by a doctoral scholarship from the École Normale Supérieure de Lyon. RPR acknowledges funding from the Swiss National Science Foundation, Project grant n° 31003A_182386. Authors would like to thank Sylvain Billiard and two anonymous reviewers for insightful comments on the present manuscript.

Version 5 of this preprint has been peer-reviewed and recommended by Peer Community In Ecology (https://doi.org/10.24072/pci.ecology.100089).

## Supplementary material

An appendix containing extra mathematical precisions and figure is available online: https://www.biorxiv.org/content/10.1101/570580v5.supplementary-material

## References

1. Ferriere R, Legendre S. 2013 Eco-evolutionary feedbacks, adaptive dynamics and evolutionary rescue theory. Phil Trans R Soc B 368, 20120081. (doi:10.1098/rstb.2012.0081)

2. Carlson SM, Cunningham CJ, Westley PAH. 2014 Evolutionary rescue in a changing world. Trends Ecol. Evol. 29, 521–530. (doi:10.1016/j.tree.2014.06.005)

3. Toby Kiers E, Palmer TM, Ives AR, Bruno JF, Bronstein JL. 2010 Mutualisms in a changing world: an evolutionary perspective. Ecol. Lett. 13, 1459–1474. (doi:10.1111/j.1461-0248.2010.01538.x)

4. Willmer P. 2011 Pollination and Floral Ecology. Princeton University Press.

5. Jordano P, Forget P-M, Lambert JE, Böhning-Gaese K, Traveset A, Wright SJ. 2010 Frugivores and seed dispersal: mechanisms and consequences for biodiversity of a key ecological interaction. Biol. Lett., rsbl20100986. (doi:10.1098/rsbl.2010.0986)

6. Wilson GWT, Rice CW, Rillig MC, Springer A, Hartnett DC. 2009 Soil aggregation and carbon sequestration are tightly correlated with the abundance of arbuscular mycorrhizal fungi: results from long-term field experiments. Ecol. Lett. 12, 452–461. (doi:10.1111/j.1461-0248.2009.01303.x)

7. Potts SG, Biesmeijer JC, Kremen C, Neumann P, Schweiger O, Kunin WE. 2010 Global pollinator declines: trends, impacts and drivers. Trends Ecol. Evol. 25, 345–353. (doi:10.1016/j.tree.2010.01.007)

8. Tylianakis JM, Didham RK, Bascompte J, Wardle DA. 2008 Global change and species interactions in terrestrial ecosystems. Ecol. Lett. 11, 1351–1363. (doi:10.1111/j.1461-0248.2008.01250.x)

9. Loeuille N. 2019 Eco-evolutionary dynamics in a disturbed world: implications for the maintenance of ecological networks. F1000Research 8. (doi:10.12688/f1000research.15629.1)

10. Parvinen K. 2005 Evolutionary suicide. Acta Biotheor. 53, 241–264. (doi:10.1007/s10441-005-2531-5)

11. Darwin C. 1877 On the various contrivances by which British and foreign orchids are fertilised by insects.

12. Parmesan C. 2006 Ecological and Evolutionary Responses to Recent Climate Change. Annu. Rev. Ecol. Evol. Syst. 37, 637–669. (doi:10.1146/annurev.ecolsys.37.091305.110100)

13. Hopkins R, Rausher MD. 2012 Pollinator-Mediated Selection on Flower Color Allele Drives Reinforcement. Science 335, 1090–1092. (doi:10.1126/science.1215198)

14. Bodbyl Roels SA, Kelly JK. 2011 Rapid Evolution Caused by Pollinator Loss in Mimulus Guttatus. Evolution 65, 2541–2552. (doi:10.1111/j.1558-5646.2011.01326.x)

15. Gervasi DDL, Schiestl FP. 2017 Real-time divergent evolution in plants driven by pollinators. Nat. Commun. 8, 14691. (doi:10.1038/ncomms14691)

16. Johnson NC, Rowland DL, Corkidi L, Egerton-Warburton LM, Allen EB. 2003 Nitrogen Enrichment Alters Mycorrhizal Allocation at Five Mesic to Semiarid Grasslands. Ecology 84, 1895–1908. (doi:10.1890/0012-9658(2003)084[1895:NEAMAA]2.0.CO;2)

17. Egerton-Warburton LM, Johnson NC, Allen EB. 2007 Mycorrhizal Community Dynamics Following Nitrogen Fertilization: A Cross-Site Test in Five Grasslands. Ecol. Monogr. 77, 527–544. (doi:10.1890/06-1772.1)

18. Nijjer S, Rogers WE, Lee C-TA, Siemann E. 2008 The effects of soil biota and fertilization on the success of Sapium sebiferum. Appl. Soil Ecol. 38, 1–11. (doi:10.1016/j.apsoil.2007.08.002)

19. Thébault E, Fontaine C. 2010 Stability of Ecological Communities and the Architecture of Mutualistic and Trophic Networks. Science 329, 853–856. (doi:10.1126/science.1188321)

20. Rohr RP, Saavedra S, Bascompte J. 2014 On the structural stability of mutualistic systems. Science 345, 1253497. (doi:10.1126/science.1253497)

21. Saavedra S, Rohr RP, Olesen JM, Bascompte J. 2016 Nested species interactions promote feasibility over stability during the assembly of a pollinator community. Ecol. Evol. 6, 997–1007. (doi:10.1002/ece3.1930)

22. Goh BS. 1979 Stability in Models of Mutualism. Am. Nat. 113, 261–275.

23. Ferriere R, Bronstein JL, Rinaldi S, Law R, Gauduchon M. 2002 Cheating and the evolutionary stability of mutualisms. Proc. R. Soc. Lond. B Biol. Sci. 269, 773–780. (doi:10.1098/rspb.2001.1900)

24. De Mazancourt C, Schwartz MW. 2010 A resource ratio theory of cooperation. Ecol. Lett. 13, 349–359. (doi:10.1111/j.1461-0248.2009.01431.x)

25. Georgelin E, Loeuille N. 2016 Evolutionary response of plant interaction traits to nutrient enrichment modifies the assembly and structure of antagonistic-mutualistic communities. J. Ecol. 104, 193–205. (doi:10.1111/1365-2745.12485)

26. Valdovinos FS, Brosi BJ, Briggs HM, Moisset de Espanés P, Ramos-Jiliberto R, Martinez ND. 2016 Niche partitioning due to adaptive foraging reverses effects of nestedness and connectance on pollination network stability. Ecol. Lett. 19, 1277–1286. (doi:10.1111/ele.12664)

27. Cheptou P, Massol F. 2009 Pollination Fluctuations Drive Evolutionary Syndromes Linking Dispersal and Mating System. Am. Nat. 174, 46–55. (doi:10.1086/599303)

28. Lepers C, Dufay M, Billiard S. 2014 How does pollination mutualism affect the evolution of prior self-fertilization? A model. Evolution 68, 3581–3598. (doi:10.1111/evo.12533)

29. Astegiano J, Massol F, Vidal MM, Cheptou P-O, Jr PRG. 2015 The Robustness of Plant-Pollinator Assemblages: Linking Plant Interaction Patterns and Sensitivity to Pollinator Loss. PLOS ONE 10, e0117243. (doi:10.1371/journal.pone.0117243)

30. Thomann M, Imbert E, Devaux C, Cheptou P-O. 2013 Flowering plants under global pollinator decline. Trends Plant Sci. 18, 353–359. (doi:10.1016/j.tplants.2013.04.002)

31. Goh BS. 1976 Global stability in two species interactions. J. Math. Biol. 3, 313–318. (doi:10.1007/BF00275063)

32. Holland JN, DeAngelis DL. 2010 A consumer–resource approach to the density-dependent population dynamics of mutualism. Ecology 91, 1286–1295. (doi:10.1890/09-1163.1)

33. Willmer P. 2011 Rewards and costs: the environmental economics of pollination. In Pollination and Floral Ecology, pp. 234–257. Princeton University Press.

34. Obeso JR. 2002 The costs of reproduction in plants. New Phytol. 155, 321–348. (doi:10.1046/j.1469-8137.2002.00477.x)

35. Dieckmann U, Law R. 1996 The dynamical theory of coevolution: a derivation from stochastic ecological processes. J. Math. Biol. 34, 579–612. (doi:10.1007/BF02409751)

36. Geritz S a. H, Kisdi E, Mesze NA G, Metz J a. J. 1998 Evolutionarily singular strategies and the adaptive growth and branching of the evolutionary tree. Evol. Ecol. 12, 35–57. (doi:10.1023/A:1006554906681)

37. Christiansen FB. 1991 On Conditions for Evolutionary Stability for a Continuously Varying Character. Am. Nat. 138, 37–50.

38. Marrow P, Dieckmann U, Law R. 1996 Evolutionary dynamics of predator-prey systems: an ecological perspective. J. Math. Biol. 34, 556–578. (doi:10.1007/BF02409750)

39. Dieckmann U, Ferrière R. 2004 Adaptive dynamics and evolving biodiversity. In Evolutionary conservation biology, pp. 188–224. Cambridge: Cambridge University Press.

40. Fishman MA, Hadany L. 2010 Plant–pollinator population dynamics. Theor. Popul. Biol. 78, 270–277. (doi:10.1016/j.tpb.2010.08.002)

41. Reekie E, Bazzaz FA. 2011 Reproductive Allocation in Plants. Elsevier.

42. Dercole F, Ferriere R, Rinaldi S. 2002 Ecological Bistability and Evolutionary Reversals Under Asymmetrical Competition. Evolution 56, 1081–1090. (doi:10.1111/j.0014-3820.2002.tb01422.x)

43. Scheffer M, Carpenter SR. 2003 Catastrophic regime shifts in ecosystems: linking theory to observation. Trends Ecol. Evol. 18, 648–656. (doi:10.1016/j.tree.2003.09.002)

44. Dieckmann U, Marrow P, Law R. 1995 Evolutionary cycling in predator-prey interactions: population dynamics and the red queen. J. Theor. Biol. 176, 91–102. (doi:10.1006/jtbi.1995.0179)

45. Biesmeijer JC et al. 2006 Parallel Declines in Pollinators and Insect-Pollinated Plants in Britain and the Netherlands. Science 313, 351–354. (doi:10.1126/science.1127863)

