## Supplementary material for "Eco-evolutionary dynamics further weakens mutualistic interaction and coexistence under population decline"

### Electronic Supplementary material

#### A) The adaptive dynamics method

In this part the symbol $*$ signal the ecological equilibrium and $\hat{\square}$ the evolutionary one.

##### The Canonical Equation

Adaptive dynamics rest on a series of hypotheses. This method models explicitly the evolutionary consequences on species density dynamics, and the feedback of species density on the evolutionary process [1,2]. Evolution occurs via small mutation steps between which plant and pollinator densities reach the ecological equilibrium. Adaptive dynamics also assume clonal reproduction and a small phenotypic impact of the mutations. The differential equation describing the evolution of the phenotypic traits, known as the canonical equation [1], is given by:

$$\frac{d\alpha}{dt}=\frac{1}{2}\mu\sigma^2 P^*(\alpha)\left.\frac{\partial\omega(\alpha_m,\alpha)}{\partial\alpha_m}\right|_{\alpha_m\to\alpha} \qquad \text{(A1)}$$

As explained in the main text the term $\frac{1}{2}\mu\sigma^2 P^*(\alpha)$ corresponds to the phenotypic variability brought by the mutation process; with $\mu$ the per individual mutation rate, $\sigma^2$ the variance of the mutation phenotypic effect, and $P^*(\alpha)$ the plant equilibrium density. The last term is the selective gradient and represents the effects of natural selection, via variations of the relative fitness of mutants $\alpha_m$ given a resident population of attractiveness $\alpha$. Therefore, the sign of the selective gradient gives the direction of evolution; a positive gradient selects larger attractiveness, while a negative gradient selects smaller trait values. The relative fitness of a mutant at a very low density is explicitly derived from the ecological dynamics (equation (1) in main text). It is computed as the *per capita* growth rate of a rare mutant population in a resident population at ecological equilibrium :

$$\omega(\alpha_m,\alpha)=\frac{1}{P_m}\frac{dP_m}{dt}\bigg|_{P_m\to 0}=r_P(\alpha_m)-c_P P^*(\alpha)+\alpha_m\gamma_A A^*(\alpha), \tag{A2}$$

with $P_m$ the mutant population density, $P^*(\alpha)$ and $A^*(\alpha)$ given by equation (3) in the main text. Remember that, due to the allocation costs, the plant intrinsic growth rate varies with the level of attractiveness $r_P(\alpha)$ .

#### The singular strategies

Eco-evolutionary dynamics (equation A1) may exhibit equilibrium points, called evolutionary singular strategies. They correspond to trait values at which equation (A1) is at equilibrium, i.e., trait variation goes to zero.

Trait variations are given by the Canonical equation (A1). Because the first part of this equation is positive, the direction of trait variations is entirely determined by the selection gradient. When it is positive, higher trait values are selected, while negative selection gradients lead to smaller traits. Here the selection gradient corresponds to the slope of the fitness function A2 at the resident trait $\alpha$ , given a small variation in the trait $\alpha_m$ .

$$\frac{\partial\omega(\alpha_m,\alpha)}{\partial\alpha_m}=\frac{dr_P(\alpha_m)}{d\alpha_m}+\gamma_A A^*(\alpha), \tag{A3}$$

Because of the hypothesis of small mutations, this yields:

$$\frac{\partial\omega(\alpha_m,\alpha)}{\partial\alpha_m}\bigg|_{\alpha_m\to\alpha}=\frac{dr_P(\alpha_m)}{d\alpha_m}\bigg|_{\alpha_m\to\alpha}+\gamma_A A^*(\alpha), \tag{A4}$$

Because all other terms of the Canonical Equation (A1) are positive, the evolutionary singular strategy ( $\hat{\alpha}$ ) correspond to trait values at which the selection gradient is null:

$$\frac{\partial\omega(\alpha_m,\alpha)}{\partial\alpha_m}\bigg|_{\alpha_m,\alpha\to\hat{\alpha}}=\frac{dr_P(\alpha_m)}{d\alpha_m}\bigg|_{\alpha_m,\alpha\to\hat{\alpha}}+\gamma_A A^*(\hat{\alpha})=0, \tag{A5}$$

with $r_P(\alpha)$ defined by equation (6) and $A^*$ by equation (3) in the main article.

This means that a singularity is obtained only when costs in terms of energy of alternative means of reproduction ( $\left.\frac{\partial r_P(\alpha_m)}{\partial \alpha_m}\right|_{\alpha_m,\alpha \to \hat{\alpha}}$ ) match the benefits in terms of pollination of increased attractiveness ( $\gamma_A A^*(\hat{\alpha})$ ).

Replacing $r_P$ , we obtain:

$$\left.\frac{\partial \omega(\alpha_m,\alpha)}{\partial \alpha_m}\right|_{\alpha_m,\alpha \to \hat{\alpha}} = \frac{-\left(\frac{\hat{\alpha}}{\alpha_{max}}\right)^s \left(1-\left(\frac{\hat{\alpha}}{\alpha_{max}}\right)^s\right)^{\frac{1}{s}-1}}{\hat{\alpha}} + \gamma_A \frac{\hat{\alpha}\gamma_P r_P(\hat{\alpha}) + c_P r_A}{c_A c_P - \hat{\alpha}^2 \gamma_A \gamma_P} = 0, \quad \text{(A6)}$$

In the linear case (i.e. when $s=1$ ), the singular strategy formula is:

$$\hat{\alpha} = \frac{c_P(c_A - \alpha_{max}\gamma_A r_A)}{\alpha_{max}\gamma_A \gamma_P} \quad \text{(A7)}$$

This solution is feasible (i.e. positive and in a plausible range value), with $\alpha_{max} < \alpha_{cl}$ as defined in equation (4) of the main text, if and only if $0 < c_A < \alpha_{max}\gamma_A r_A$ ; i.e. the intraspecific competitive losses need to stay below the maximal energetic gain of the animal.

In this linear case, increasing plant or animal losses or decreasing animal intrinsic growth rate (within the conditions for a feasible solution) will increase the singular strategy value, meaning that when these losses are high or the animal intrinsic growth low the plant invest more in the mutualistic interaction. On the contrary higher gains from the mutualistic interaction will lead into a lower investment in the plant attractiveness at eco-evolutionary equilibrium.

Aside from its existence, a singular strategy can be an endpoint of evolution if convergent (evolutionary dynamics locally lead to it) and non-invasible (it persists in time because of a resistance to invasion by nearby mutants).The mathematical computation for the existence of singular strategies and their convergence and invasibility properties are given in the following part.

### B) Convergence and invasibility properties of the singular strategies

#### The conditions for invasibility

With the trade-off function defined in main text equation (5) we can differentiate the fitness function a second time to analyse the convergence and invasibility of the singular strategies, to deduce the overall trait dynamics [1]. The singular strategy ( $\hat{\alpha}$ ) is non-invasible (ie, an ESS [5]) when:

$$\left.\frac{\partial\omega^2(\alpha_m,\alpha)}{(\partial\alpha_m)^2}\right|_{\alpha_m,\alpha\rightarrow\hat{\alpha}}=\frac{(1-s)\left(\frac{\hat{\alpha}}{\alpha_{max}}\right)^s\left(1-\left(\frac{\hat{\alpha}}{\alpha_{max}}\right)^s\right)^{\frac{1}{s}-2}}{\hat{\alpha}^2}<0 \qquad \text{(B8)}$$

Concave trade-offs ( $s>1$ ) therefore lead to non-invasible singular strategies, while convex trade-offs ( $s<1$ ) yield invasible strategies.

In the case of a linear trade-off equation (B8) is equal to 0, the strategy is neutral from an invasibility point of view.

#### The conditions for convergence

The previous equation, summed with the crossed derivation of the fitness function gives conditions for convergence of the singular strategy [1]. The singular strategy is convergent when:

$$\left.\frac{\partial\omega^2(\alpha_m,\alpha)}{(\partial\alpha_m)^2}\right|_{\alpha_m,\alpha\rightarrow\hat{\alpha}}+\left.\frac{\partial\omega^2(\alpha_m,\alpha)}{\partial\alpha\,\partial\alpha_m}\right|_{\alpha_m,\alpha\rightarrow\hat{\alpha}}<0 \qquad \text{(B9)}$$

The above-mentioned formula requires the calculation of the cross-derivation. Using results from equation (A3) , and with small mutation close to the singular strategy ( $\alpha_m,\alpha\rightarrow\hat{\alpha}$ ), it gives:

$$\left.\frac{\partial \omega^2(\alpha_m,\alpha)}{\partial \alpha \partial \alpha_m}\right|_{\alpha_m,\alpha \to \hat{\alpha}} = \gamma_A \frac{d A^*(\hat{\alpha})}{d \alpha} \qquad \text{(B10)}$$

According to the formula of $A^*(\hat{\alpha})$ given in equation (3) of the main article, the previous equation is equivalent to:

$$\left.\frac{\partial \omega^2(\alpha_m,\alpha)}{\partial \alpha \partial \alpha_m}\right|_{\alpha_m,\alpha \to \hat{\alpha}} = \frac{\gamma_A \gamma_P (2 c_P \gamma_A r_A \hat{\alpha} + (c_A c_P + \hat{\alpha}^2 \gamma_A \gamma_P) r_P(\hat{\alpha}) + \hat{\alpha}(c_A c_P - \hat{\alpha}^2 \gamma_A \gamma_P) r_P'(\hat{\alpha})}{(c_A c_P - \hat{\alpha}^2 \gamma_A \gamma_P)^2} \qquad \text{(B11)}$$

with $r_P(\hat{\alpha})$ defined by equation (6) in the main article, and $r_P'(\hat{\alpha}) = r_P(\hat{\alpha}) \frac{1}{\hat{\alpha}\left(1 - \left(\frac{\hat{\alpha}}{\alpha_{max}}\right)\right)^{-s}}$ .

The sum of equation (B11) at the eco-evolutionary equilibrium (i.e. when $\alpha_{max}, \alpha \to \hat{\alpha}$ ) and (B8) is however too complex in the general case to give a simple to understand the convergence condition (as required by equation B9)

In the linear case (i.e. when $s=1$ ), equation (B11) at the eco-evolutionary equilibrium becomes:

$$\left.\frac{\partial \omega^2(\alpha_m,\alpha)}{\partial \alpha \partial \alpha_m}\right|_{\alpha_m,\alpha \to \hat{\alpha}} = \frac{\gamma_A \gamma_P \left(c_A c_P + \hat{\alpha}^2 \gamma_A \gamma_P + 2 c_P \hat{\alpha}\left(\gamma_A r_A - \frac{c_A}{\alpha_{max}}\right)\right)}{(c_A c_P - \hat{\alpha}^2 \gamma_A \gamma_P)^2} \qquad \text{(B12)}$$

Because then equation (B8) $\left.\frac{\partial \omega^2(\alpha_m,\alpha)}{(\partial \alpha_m)^2}\right|_{\alpha_m,\alpha \to \hat{\alpha}} = 0$ , the convergence condition then depends only on the above cross derivation (B12).

According to equation (A7), $\gamma_A r_A - \frac{c_A}{\alpha_{max}} = \frac{-\hat{\alpha} \gamma_A \gamma_P}{c_P}$ , equation (B12) when s=1 is equal to:

$$\left.\frac{\partial \omega^2(\alpha_m,\alpha)}{\partial \alpha \partial \alpha_m}\right|_{\alpha_m,\alpha \to \hat{\alpha}} = \frac{\gamma_A \gamma_P}{(c_A c_P - \hat{\alpha} \gamma_A \gamma_P)^2}\left[c_A c_P + \hat{\alpha}^2 \gamma_A \gamma_P - 2 c_P \hat{\alpha} \frac{\hat{\alpha} \gamma_A \gamma_P}{c_P}\right] \qquad \text{(B13)}$$

Which, when simplifying gives:

$$\left.\frac{\partial\omega^2(\alpha_m,\alpha)}{\partial\alpha\,\partial\alpha_m}\right|_{\alpha_m,\alpha\rightarrow\hat{\alpha}}=\frac{\gamma_A^2\gamma_P^2}{(c_A c_P-\hat{\alpha}\gamma_A\gamma_P)^2}-\left[\frac{c_A c_P}{\gamma_A\gamma_P}-\hat{\alpha}^2\right] \tag{B14}$$

Note that $\frac{c_A c_P}{\gamma_A\gamma_P}=\alpha_{cl}^2$ . Because $\hat{\alpha}<\alpha_{max}<\alpha_{cl}$ , when $\hat{\alpha}$ exist, the above derivation is always positive, meaning that a linear trade-off always leads to a divergent singular strategy.

#### C) Effect of the trade-off shape on the number of singular strategies

We prove in this section that $s=2$ (concave trade-off) is a threshold for the existence of a second singular strategy in the case $r_A=0$ .

For $r_A=0$ the singular strategies are the solution of the following equation (obtained by setting $r_A=0$ in equation (A6) ):

$$\left.\frac{\partial\omega(\alpha_m,\alpha)}{\partial\alpha_m}\right|_{\alpha_m,\alpha\rightarrow\hat{\alpha},r_A=0}=\frac{r_P(\hat{\alpha})^{(1-s)}\left(\hat{\alpha}^2\gamma_A\gamma_P-\left(\frac{\hat{\alpha}}{\alpha_{max}}\right)^s c_A c_P\right)}{\hat{\alpha}\left(c_A c_P-\hat{\alpha}^2\gamma_A\gamma_P\right)}=0 \tag{C15}$$

If $s>1$ , then $\hat{\alpha}=0$ is always a solution. Now we can study the existence of a second non-zero singular strategy, which is then the solution of:

$$\hat{\alpha}^2\gamma_A\gamma_P=\left(\frac{\hat{\alpha}}{\alpha_{max}}\right)^s c_A c_P \tag{C16}$$

The solution can be explored geometrically, but we first need to rewrite the equation as follow:

$$\hat{\alpha}^2=\left(\frac{\hat{\alpha}}{\alpha_{max}}\right)^s\frac{c_A c_P}{\gamma_A\gamma_P}=\left(\frac{\hat{\alpha}}{\alpha_{max}}\right)^s\alpha_{cl}^2 \tag{C17}$$

Remember that the stability conditions impose that $\frac{c_A c_P}{\gamma_A\gamma_P}=\alpha_{cl}^2>\alpha_{max}^2$ . Now, we can plot the left and the right side of equation (C17) as a function of $\hat{\alpha}$ and the solution is given at the crossing of the two curves (figure C1). Note that only the right side depends on the value of the trade-off parameter $s$. Figure C1, shows that if $1<s\leqslant 2$ there exists a unique solution that is $\hat{\alpha}=0$. This is because the right side is always larger than the left side for $\hat{\alpha}>0$. Now, if $s>2$, there exists a second solution, which is $\hat{\alpha}=\sqrt[s-2]{\frac{\alpha_{cl}^2}{\alpha_{max}^s}}$. This proof that at the value $2=s$ there is a branching from one to two singular strategies. Now, the nature of the two strategies needs to be explored numerically, as well as how it extends to negative values of $r_A<0$.

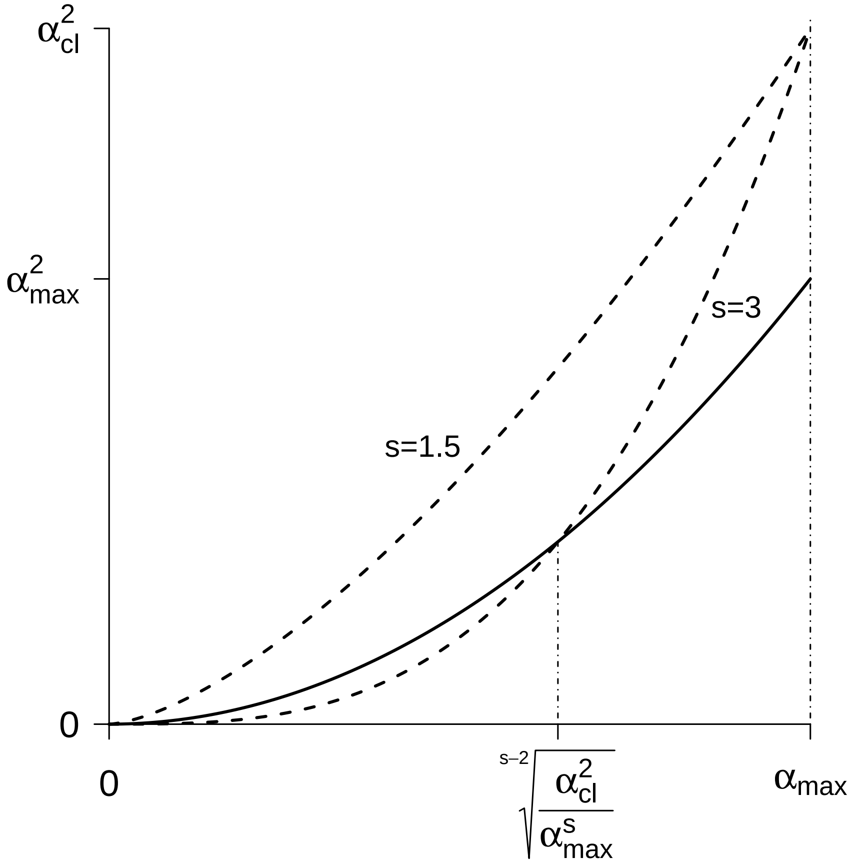


Figure C1: geometry representation of equation (C17). The black line represents the left side of equation (C17), while the right side is given by the dashed line for two different values of the trade-off shape parameters $s$.

### D) Supplementary figures

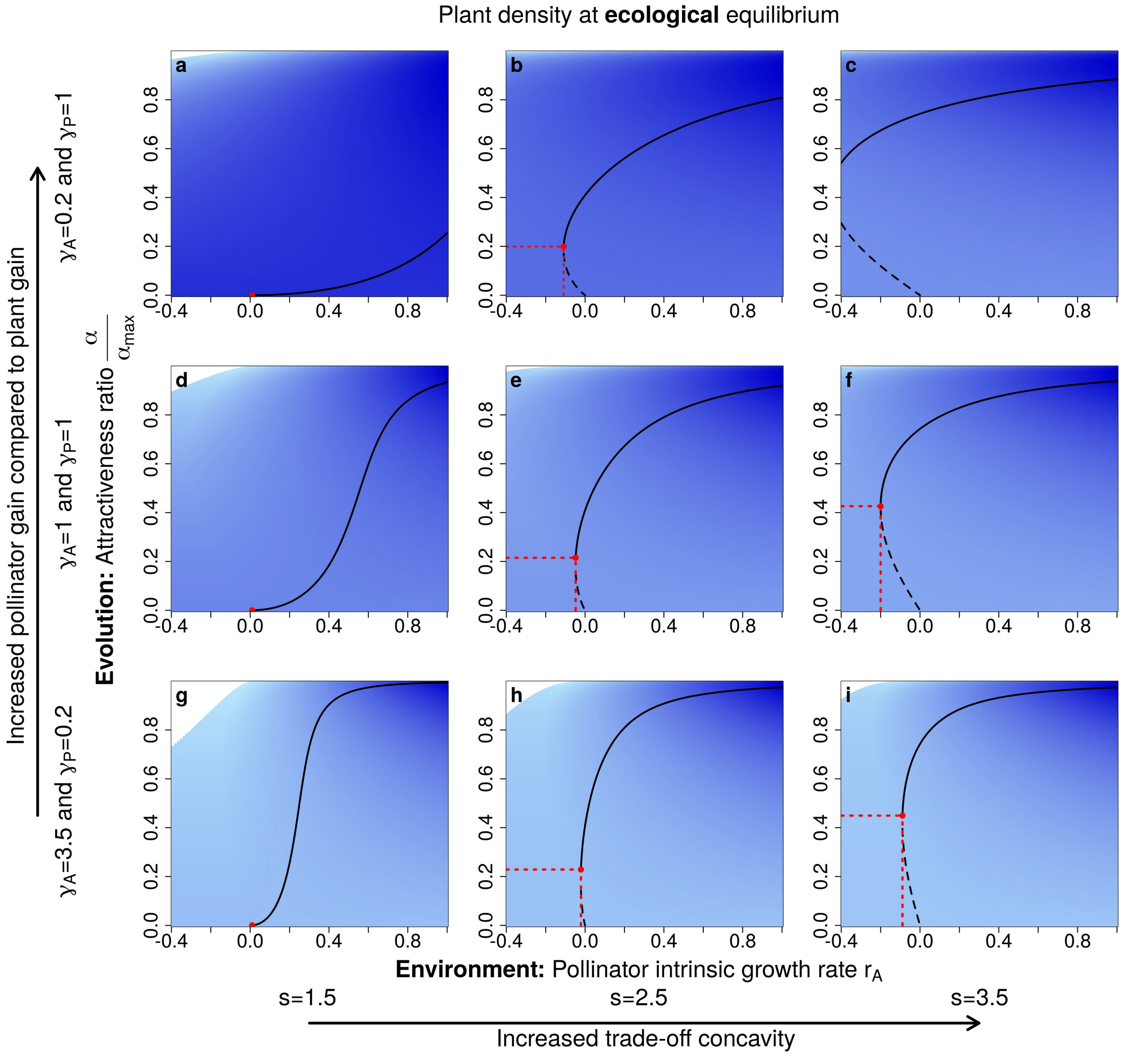


Figure D1: **Influence of trade-off shape and mutualistic gains on $E^3$ diagrams**. As in figure 5 in the main text, columns differ in trade-off concavity. Lines differ in the asymmetry of mutualistic gains: in the top line (panels a,b, and c) pollinators benefit more than plants; the middle line (panels d,e, and f) shows equal gains while in the bottom line plant gains are larger. Red points and dotted lines represent the lowest $r_A$ and $\frac{\alpha}{\alpha_{max}}$ values for maintaining a CSS, allowing the maintenance of the mutualistic interaction. Colours and lines are the same as in figure 4 in the main text. The parameter values are $c_A = c_P = 1$ and $\alpha_{max} = 0.8 * \alpha_{cl}$ .

### E) References

1. Dieckmann U, Law R. 1996 The dynamical theory of coevolution: a derivation from stochastic ecological processes. *J. Math. Biol.* **34**, 579–612. (doi:10.1007/BF02409751)

2. Geritz S a. H, Kisdi E, Mesze NA G, Metz J a. J. 1998 Evolutionarily singular strategies and the adaptive growth and branching of the evolutionary tree. *Evol. Ecol.* **12**, 35–57. (doi:10.1023/A:1006554906681)

3. Christiansen FB. 1991 On Conditions for Evolutionary Stability for a Continuously Varying Character. *Am. Nat.* **138**, 37–50.

4. Marrow P, Dieckmann U, Law R. 1996 Evolutionary dynamics of predator-prey systems: an ecological perspective. *J. Math. Biol.* **34**, 556–578. (doi:10.1007/BF02409750)

5. Maynard Smith J. 1982 *Evolution and the Theory of Games*. Cambridge University Press.
